# Characterization of expression quantitative trait loci in extensively phenotyped pedigrees ascertained for bipolar disorder

**DOI:** 10.1101/031427

**Authors:** C. Peterson, S. Service, A. Jasinska, F. Gao, I. Zelaya, T. Teshiba, C. Bearden, V. Reus, G. Macaya, C. López-Jaramillo, M. Bogomolov, Y. Benjamini, E. Eskin, G. Coppola, N. Freimer, C. Sabatti

## Abstract

The observation that variants regulating gene expression (expression quantitative trait loci, eQTL) are at a high frequency among SNPs associated with complex traits has made the genome-wide characterization of gene expression an important tool in genetic mapping studies of such traits. As part of a study to identify genetic loci contributing to bipolar disorder and a wide range of BP-related quantitative traits in members of 26 pedigrees from Costa Rica and Colombia, we measured gene expression in lymphoblastoid cell lines derived from 786 pedigree members. The study design enabled us to comprehensively reconstruct the genetic regulatory network in these families, provide estimates of heritability, identify eQTL, evaluate missing heritability for the eQTL, and quantify the number of different alleles contributing to any given locus.

## INTRODUCTION

Dozens of investigations have now shown that the identification of local eQTL may play a crucial role in delineating the causal variant(s) contributing to genetic associations observed for complex disorders or quantitative traits. While it may be particularly informative to evaluate, for a given trait, eQTL specific to tissues implicated in the manifestation of that trait, this strategy may be infeasible for human brain related traits, such as psychiatric disorders and their endophenotypes. In this study we report the results of gene expression in lymphoblastoid cell lines (LCL) for 786 genotyped members of Costa Rican and Colombian pedigrees investigated for severe bipolar disorder (BP1) and quantitative phenotypes from the domains of temperament, neurocognition, neuroanatomy, sleep, and circadian activity (Fears et al. 2014; Pagani et al., in press). We selected LCL for ease of study and on the basis of the increasing evidence that a substantial proportion of local genetic regulation is conserved across tissues (Ding et al. 2010; Nica et al. 2011; Flutre et al. 2013). Studying LCLs has enabled at least a partial reconstruction of the specific regulatory network of these families, allowing us to identify those components that might show differences from the general population. We study the genetic regulation of expression in these pedigrees at a multiscale level: we estimate heritability, evaluate the relative importance of local vs. distal genomic variation, identify variants with regulatory effects, and analyze the role of multiple associated SNPs in the same region. By capitalizing on known pedigree structure, as well as extensive genotyping, we can compare different methodologies for heritability estimation. The most interesting element of regulatory networks for our purpose is the localization of SNPs with regulatory effects (eSNPs): these variants are candidates for association to the BP1 endophenotypes measured in our sample. To control the rate of false discoveries of eSNPs, we adopt a novel hierarchical testing procedure that leads to the analysis of expression quantitative trait loci (eQTL) data in a stage-wise manner with increasing levels of detail.

## METHODS

### Sample collection

The study subjects are members of 26 Costa Rican and Colombian pedigrees ascertained from local hospitals and clinics based on multiple individuals affected with BP1. Descriptions of pedigrees and ascertainment procedures are provided in Fears et al. (2014). Written informed consent was obtained from each participant, and institutional review boards at participating sites approved all study procedures.

### RNA Extraction and Measurement of Gene Expression

Lymphoblastoid cell lines (LCLs) were established at two sites, and RNA was extracted from these cell lines and its expression quantified using Illumina Human HT-12 v4.0 Expression BeadChips. Expression values were background corrected, quantile normalized, log2 transformed, and corrected for major known batch effects. The outcome of these procedures is what we refer to as ‘probe expression’ for all subsequent analyses. After quality control filters, the 34,030 probes included in the final set were uniquely aligned to hg19, contained no common SNPs (as defined in dbSNP 137 or 138), queried 24,385 unique genes, and their expression was detected in at least one individual. For a detailed description of the processing steps used at each site and the RNA quantification, normalization, and quality control procedures, see the supplementary note.

### DNA Extraction, Genotyping, and Subject Inclusion Criteria

DNA was extracted from blood or LCLs using standard protocols. Illumina Omni 2.5 chips were used for genotyping, in three batches. A subset of samples was repeated in each batch to enable concordance checks. A total of 2,026,257 SNPs were polymorphic and passed all QC procedures, including the evaluation of call rate, testing for Hardy Weinberg equilibrium, and Mendelian error. A total of 1,024,051 autosomal SNPs with MAF of at least 10% were selected for use in the subsequent association analysis. After excluding married-ins with no descendants in the study and cases of possible contamination, the analyzed sample contains 786 individuals with both genotype and gene expression data. (See supplementary note for details.)

### Adjustment for factors affecting global gene expression

In order to adjust for both known and unknown factors affecting global gene expression, all association and heritability analyses include age, sex and batch as covariates, in addition to a set of PEER factors to adjust for latent determinants of global gene expression (Stegle et al. 2012). We chose to include 20 PEER factors on the basis of the proportion of global gene expression explained, and found that these PEER factors were strongly correlated with batch, but not with family groupings, suggesting that they are in fact correcting for technical artifacts.

### Relationship between Gene Expression Levels and BP1

We focus here on two categories of related individuals: those with a clinical diagnosis of BP1 vs. those with no history of BP1. We would like to note that the subjects with no history of BP1 are not necessarily healthy controls; they are included in the sample because they are related to a BP1 subject and may have a range of other non-BP1 diagnoses. To identify genes with differential expression in the BP1 vs. non-BP1 individuals accounting for the correlation induced by relatedness, we used a variance components approach, as implemented in Mendel (Lange et al. 2013). Specifically, we computed p-values for the association of BP1 to gene expression by fitting a variance components model with additive genetic and environmental components and BP1 status as predictor, as well as additional covariates corresponding to age, sex, batch and PEER factors. To correct for the multiplicity of tests, we applied the Benjamini-Hochberg (BH) procedure (Benjamini and Hochberg 1995) to control the false discovery rate (FDR) to the 5% level.

### Local vs. Distal Genetic Regulation

The eQTL literature documents a distinction between cis vs. trans regulation, although the precise definition of these is sometimes elusive. Following the suggestion of Albert and Kruglyak (2015), we adopt the terminology “local” and “distal” regulation to distinguish the situations where genetic variants and the genes whose expression they regulate are nearby or far away in the genome, without any assumption on the mechanisms of this regulation. Operationally, we define “local” associations as those between SNPs and probes where the SNP is located within 1Mb of either end of the probe, and “distal” as all other probe-SNP associations, including those across different chromosomes.

### Heritability of Gene Expression

For each probe, we estimated the heritability of gene expression using two approaches: a variance components model relying on known family relationships as implemented in Mendel (Lange et al. 2013), and a variance decomposition based on observed genotypic similarities among individuals as implemented in GCTA (Yang et al. 2011). Both analyses included age, sex, batch and PEER factors as covariates. In our primary GCTA analysis, we utilized a genetic relatedness matrix (GRM) based on the full set of genome-wide SNPs. This allowed us to calculate the ratio of genetic variability over total phenotypic variability for each probe. We then compared the estimates obtained using Mendel and GCTA. To determine which probes were significantly heritable, we relied on the likelihood ratio test implemented in GCTA to obtain p-values for the significance of the genetic variance component.

As a secondary analysis, we used GCTA to refine the variance decomposition of probe expression to obtain estimates of the proportion of probe heritability due to local regulation. Specifically, we utilized the multiple GRM option in GCTA with two GRMs specified: one based on the set of SNPs within 1Mb of the probe of interest (whenever a sufficient number of SNPs was present), and one based on all SNPs genome-wide (a reasonable stand-in for relatedness based on distal SNPs). This strategy allowed us to partition the heritability into local vs. global components and calculate the ratio of local genetic variability to total variability.

With regards to interpretation of the resulting estimates, we note that the goal of GCTA is to estimate the additive effects of the genotyped SNPs, rather than a true estimate of heritability. Yang et al. (2011) therefore recommend excluding related subjects since including these will bias the estimate of the proportion of variance explained by common variants upward due to factors such as shared environment or rare variants passed down within a family. Since we include related subjects, our GCTA results will be inflated relative to those for unrelated subjects, and therefore are more similar to the family-based heritability estimates.

### Computation of SNP-probe association p-values

We computed association p-values for each SNP-probe pair using the pedigree GWAS option in Mendel including additive genetic and environmental variance components (Lange et al. 2013; Zhou et al. 2014). The Mendel implementation relies on a score test to greatly increase the speed of computation of association p-values in mixed models. For the most promising SNP-probe pairs, a standard likelihood ratio test (LRT) is conducted, and effect sizes are derived. In our analysis, we included age, sex, batch and PEER factors as covariates. We performed the LRT for the 100 most significant local and 100 most significant distal SNPs for each probe, with the score test used for the remaining SNP-probe pairs.

### Multiplicity adjustment and identification of significant results

Our hierarchical testing approach is based on the selective procedure by Benjamini and Bogomolov (2014) whose effectiveness in genetic association studies for multiple phenotypes has been explored in Peterson et al. (2015a). We rely on the software implementation provided in the `TreeQTL` R package (Peterson et al. 2015b). The testing procedure is designed to take into account that local regulation is more common than distal (the hypotheses in these two classes are tested separately) and that SNPs with distal effects are likely to affect the expression of more than one probe. While the possibility of identifying variants involved in the local regulation of each probe depends on the sample size and the signal strength, it is quite reasonable to expect that the expression of every gene could be affected by appropriate sequence variation in the genomic region surrounding it. In contrast, one expects that only a small portion of the genotyped variants have any regulatory role. Both to capitalize on this heterogeneity and because our ultimate interest is to identify genetic variants that have phenotypic effects, we apply a multiscale testing strategy to first identify SNPs that have regulatory effects (eSNPs). We control the FDR in these discoveries at a target level of 0.05 with the Benjamini-Yekutieli (2001) procedure, a conservative approach which is robust to dependence among the test statistics and therefore appropriate given linkage disequilibrium among the SNPs. In a second stage we investigate which specific probes are influenced by these eSNPs. We control the expected average proportion of false SNP-probe associations across the selected SNPs at a target 0.05 level with the Benjamini Bogomolov (BB) method (Benjamini and Bogomolov 2014), which has been shown in Peterson et al. (2015a) to control the relevant error rates under the typical dependency structure of multi-trait GWAS.

### Genomic characteristics of eSNPs

We studied the position of local eSNPs relative to the transcription start site (TSS) of the gene queried by the probe to which they were associated. TSS information was derived from the UCSC Genome Browser (http://genome.ucsc.edu/). We investigated the distal eSNPs by assessing their overlap with local eSNPs and by comparing their locations with the annotations derived by the Roadmap Epigenomics Project (http://egg2.wustl.edu/roadmap/web_portal/) for LCLs using ChIP-Seq and DNAse-Seq (Roadmap Epigenomics Consortium 2015).

### Cross-study comparisons

Cross-study comparisons are hampered by many factors including changes in annotation resulting in different gene symbols, changes in SNP names, and the use of different versions of the human physical map. We downloaded results from eQTL analysis of blood or LCL from the seeQTL database (http://www.bios.unc.edu/research/genomic_software/seeQTL/), including results from Zeller et al. (2010), Wright et al. (2014), and a meta-analysis of HapMap LCLs, and also obtained results of Westra et al. (2013) for associations with FDR less than 50%. We used official gene symbols to compare results across studies.

### eSNP effect sizes, and percentage of heritability explained

For each probe associated to some of the discovered eSNPs, we constructed a multivariate linear mixed model relating expression to the genotypes at significant SNPs, local or distal. Using the variance components model implemented in Mendel, a fixed effect was estimated for age, sex, batch, the PEER factors, and each of the genetic variants, while a random effect was used to capture family structure. We then calculated the proportion of variance explained in this model by the collection of local eSNPs and distal eSNPs and compared it with the local and global heritability estimates obtained using the partitioning approach of GCTA.

To account for the fact that linkage disequilibrium may lead to the identification of a number of neighboring SNPs as associated to the same probe - even when the underlying association is effectively captured by one SNP alone - we performed model selection to determine the number of SNPs that might reasonably correspond to independent signals. Specifically, after transforming the data to obtain independent observations (using the appropriate variance covariance matrix determined from the mixed model analysis in Mendel), for each probe we carried out stepwise forward selection, relying on the BIC criteria, and using residual expression (adjusted for covariates) as the response and the eSNPs associated to the probe as the pool of predictors. This procedure gave us an estimate of the number of independent eSNPs affecting each probe, as well as the value of the percentage of variance explained (the adjusted r^2^ value) for the resulting multivariate linear model. For comparison, we also obtained the percentage of variance explained (the r^2^ value) for the univariate linear model using the most strongly associated eSNP (local or distal) as the only predictor. We then computed the ratio of the r^2^ for each model to the heritability previously estimated using the variance components model in Mendel.

## RESULTS

### Relationship between Gene Expression Levels and BP1

We did not detect statistically significant differences between the residual mean expression for BP1 subjects (n=193) and their non-BP1 relatives (n=593) after correcting for multiple comparisons. One probe had p<5e-05: ILMN_1805371 on chromosome 19 at 8.5Mb (querying the expression of ARMCX3 on chromosome X, p=1.65e-05). However, no comparisons were significant at an FDR threshold of 5%. Since BP1 status was not associated with changes in gene expression, we did not explicitly adjust for BP1 status in the remaining analysis.

### Heritability of Gene Expression

The distribution of heritability estimates across all 34,040 probes obtained using Mendel, shown at left in Figure 1, had median 0.10 (mean=0.03). Estimates of the heritability of gene expression based on kinship obtained using Mendel correlated well with estimates of the proportion of phenotypic variation due to genome-wide SNPs obtained using GCTA (r=0.99), suggesting agreement between the known pedigree structure and levels of genetic similarity in the subjects (Supplementary Figure 1); the estimates from Mendel tended to be slightly larger than those from GCTA, however. The median proportion of variance explained by genome-wide SNPs as computed by GCTA was 0.09 (mean=0.03). The likelihood ratio test for the significance of the genetic variance component in GCTA resulted in 11,911 rejections (35%) at p<0.05; 9,977 rejections (29%) at FDR threshold 0.05; and 4,126 rejections (12%) applying the Bonferroni correction to target FWER 0.05. The median proportion of variance in gene expression explained by genome-wide SNPs among probes satisfying FDR<0.05 was 0.21 (range 0.07–1.00).

**Figure 1:**
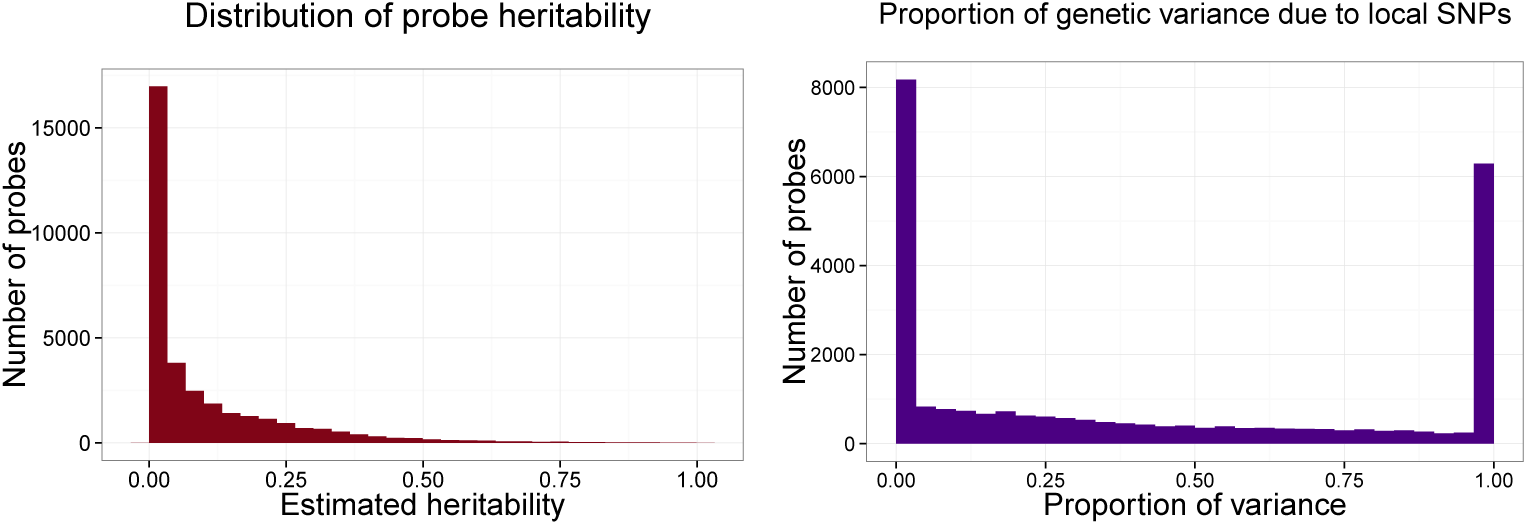
Distribution of estimated heritability of probe expression obtained using Mendel for all 34,030 probes (left), and distribution of the proportion of total genetic variance attributed to local genetic variation (right) for the 9,649 significantly heritable probes (FDR<0.05) where partitioning using the multiple GRM approach in GCTA was possible.

Among the 9,977 significantly heritable probes (FDR<0.05), 9,649 had a sufficient number of SNPs in the local region to obtain a GRM usable for partitioning; for these probes, a median of 33% of the total genetic variance was attributed to local genetic variation (mean=39%). The distribution of the proportion of total genetic variance attributed to local genetic variation for these probes is shown at right in Figure 1. Probes with a low proportion of genetic variance attributed to local genetic variation (<10%) have a significantly smaller number of local SNPs than those with a larger proportion (one-sided t-test p<2.2e-16) and are associated to a significantly higher number of distal eSNPs (one-sided t-test p=1.1e-5), suggesting that both a failure to measure relevant local SNPs and the effects of distal regulation may explain the fraction of heritable probes found to have a low local proportion of genetic variance.

### eSNP discoveries

Controlling the FDR of eSNP discoveries at a 5% level, we identify 139,668 local eSNPs and 11,016 distal eSNPs. Controlling the expected value of the average proportion of false discoveries for probe-SNP association across the discovered eSNPs to 5% as well results in the identification of 305,635 local probe-SNP pair associations and 22,304 distal probe-SNP pair associations. There are 10,065 distinct probes involved in these associations (9,645 in local regulation and 1,081 in distal, with an overlap of 661).

We now consider some of the characteristics of the discovered eSNPs. In keeping with current understanding of the mechanisms of local regulation, 72% of the local eSNPs are upstream from the gene they putatively regulate, and 15% of these are within 100kb upstream from the transcription start site (TSS). The distribution of local eSNPs by distance from the TSS, calculated as the TSS position of the queried gene minus the SNP position for each SNP-probe pair discovered, shows that the discoveries are most concentrated closest to the TSS (at left in Figure 2). Among the discovered distal eSNPs, 50% also appear to act as local regulators, a phenomenon that has been noted before (e.g. Westra et al. 2013; Bryois et al. 2014). On average, distal eSNPs affect 2.0 probes (median=1.0), or 1.8 genes; the distribution of the number of genes controlled by distal eSNPs is shown at center in Figure 2. Utilizing the annotations from the Epigenomics Roadmap, we found that 27% of distal eSNPs fall within narrow peaks (which reflect point sources such as transcription factors or chromatin marks associated with transcription start sites) and 38% fall within broad domains (which cover extended areas associated with many other types of histone modifications), indicating that a substantial portion of distal eSNPs are located within functional genomic regions. The most strongly associated local eSNP to each probe with local associations had an average effect size of magnitude 0.12; the comparable average for the distal setting was 0.21. The distributions of effect sizes for local and distal regulation are shown at right in Figure 2: the appreciable difference in effect sizes is likely due to the “winner’s curse” phenomenon given the large number of distal hypotheses. A more precise investigation of the percentage of variance explained by local and distal eSNPs is given in a later section. For a comparison of the number of discoveries under different error controlling strategies and their characteristics, see Supplementary Table 1 and Supplementary Figure 2.

**Figure 2:**
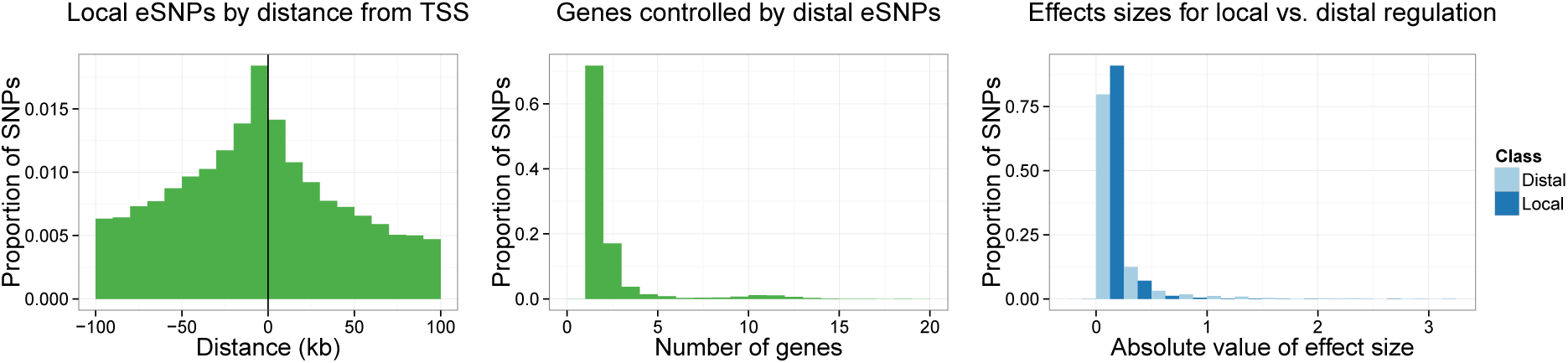
Characteristics of local and distal eSNPs. Position of local eSNPs relative to transcription start site (TSS) of the gene queried by the associated probe (left). Number of genes controlled by distal eSNPs (center), excluding SNP kgp22834062, which was associated to 129 genes. Effect sizes of the most significant SNP for each probe with any local or distal associations (right).

### Cross-study comparison of discovered eSNPs

To compare our discovered local eSNPs with those of other studies, we rely on the named genes they appear to regulate. This allows us to implicitly account both for the effect of linkage disequilibrium and the different genotypes available. Considering first local association and matching on gene name, our study and published studies had 14,174 gene names in common; 6,456 have significant local associations in our work, and 7,755 have local associations with p<0.0001 in the published studies. Of the 6,456 genes we find significant and on which we have available data in other studies, 4,790 are significant in other studies (430 are significant in one other study, 1,354 are significant in two other studies, 1,182 are significant in 3 other studies and 1,194 are significant in all 4 studies examined).

Examination of distal associations in our work and published studies indicates 10,002 gene names in common; 409 have significant distal associations in our work, and 528 genes have distal associations with p<5e-08 in the published studies. Of the 409 genes we find significant and on which we have data available in other studies, 63 are significant in other studies (17 in one other study, 24 in two studies, 16 in three studies, and 6 in all four studies examined). Only 34 of these 63 genes identified as being significantly affected by distal variants in our study were also identified as having significant distal associations in published work to SNPs on the same chromosome as ours; and 23 of these 34 genes involved associations to SNPs <2Mb apart in our study compared to the published studies (Supplementary Table 2).

We examined whether the same SNPs were involved in distal associations in multiple studies, without specifying that the associations were to the same genes. We considered this question matching both on SNP name and on SNP position, requiring that the SNPs were selected as eSNPs in our work at FDR 5% and had associations in published studies at p<5e-08. We found 33 SNPs on six chromosomes to have distal associations to one or more genes in both our study and in published studies (p<5e-08); however the distal associations were to different genes (Supplementary Table 3).

There are only ten distal associations significant in our work (controlling the expected average proportion of false associations involving the selected eSNPs to 5%) and in published studies (p<5e-08) that involve the same SNP and same gene: (1) *LIMS1* on chromosome 2 at ~10.9Mb is associated to five SNPs on chromosome 6, at 32.4–32.7Mb (rs13192471, rs3129934, rs3763313, rs9268877, rs9272219) in our work and in Westra et al.; (2) three probes in *DUSP22* on chromosome 6 at ~0.35Mb are associated to one SNP on chromosome 16 at ~35Mb (rs12447240) and is also associated to this gene in Zeller et al.; (3) *OR2AG1* on chromosome 11 at ~6.8Mb is associated to one SNP on chromosome 21 at 34.6Mb (rs1131964) in both our study and Zeller et al.; (4) *TSSC4* on chromosome 11 at ~2.4Mb is associated to one SNP on chromosome 6 at ~31.2Mb (rs3131018) in both our study and Westra et al.; (5) *NOMO1* on chromosome 16 at ~14.9Mb is associated to one SNP on chromosome 16 at ~16.3Mb (rs4780600) in both our study and in Zeller et al.; (6) and lastly *RTF1* on chromosome 15 at ~41.7Mb is associated to one SNP on chromosome 17 at 2.5Mb (rs8081803) in both our work and Zeller et al.

The sparser concordance of reconstructed distal vs. local regulation in the cross-study comparison is not surprising: the power to detect distal effects is considerably smaller in all studies, while the impact of confounders stronger. Two additional reasons might explain this difference. On the one hand, our methodology to identify distal eSNPs has larger power to discover multiple genes regulated by the same variant. On the other hand, some of our unique findings might be due to the ascertainment of the subjects, who are members of families carrying genes predisposing to BP1 and/or to extreme values of BP1-related quantitative traits.

### Proportion of heritability explained by eSNPs

To examine the explanatory power of the discovered eSNPs, we focus on the probes that they affect. Of the 10,065 probes associated to any eSNPs, 7,036 were significantly heritable at an FDR of 5% (6,770 with local associations, 903 with distal associations, and 637 with both). Among the non-heritable probes with eSNP associations, 94% had only local associations, suggesting that these discoveries reflect the less stringent multiplicity control for the discovery of local associations. When the eSNPs for each of the 7,036 heritable probes were included as fixed effects in a variance components model of the probe expression, the genetic variance component was estimated to be 0 for 1,448 (21%) of the probes, indicating that for these probes, the eSNPs capture essentially all of the genetic component of variation in probe expression. The distribution of the proportion of genetic variance due to the selected eSNPs (estimated as 1 - the ratio of the genetic variance component when eSNPs are included as fixed effects to the genetic variance component when eSNPs are not included) is shown in Figure 3, assuming values less than 0 (12%) are exactly 0. The median proportion for the set of probes with only local, only distal, or both types of associations is 0.44, 0.52, and 0.97, respectively, demonstrating that the eSNPs do explain a substantial proportion of the heritability of gene expression, particularly for probes with both significant local and significant distal associations. The distributions of the local and total genetic proportions of variance under partitioning using GCTA for probes with only local, only distal, or both types of associations (Supplementary Figure 3) demonstrates that probes with local associations do in fact have larger proportions of variance due to local effects vs. probes not associated to any local SNPs.

**Figure 3:**
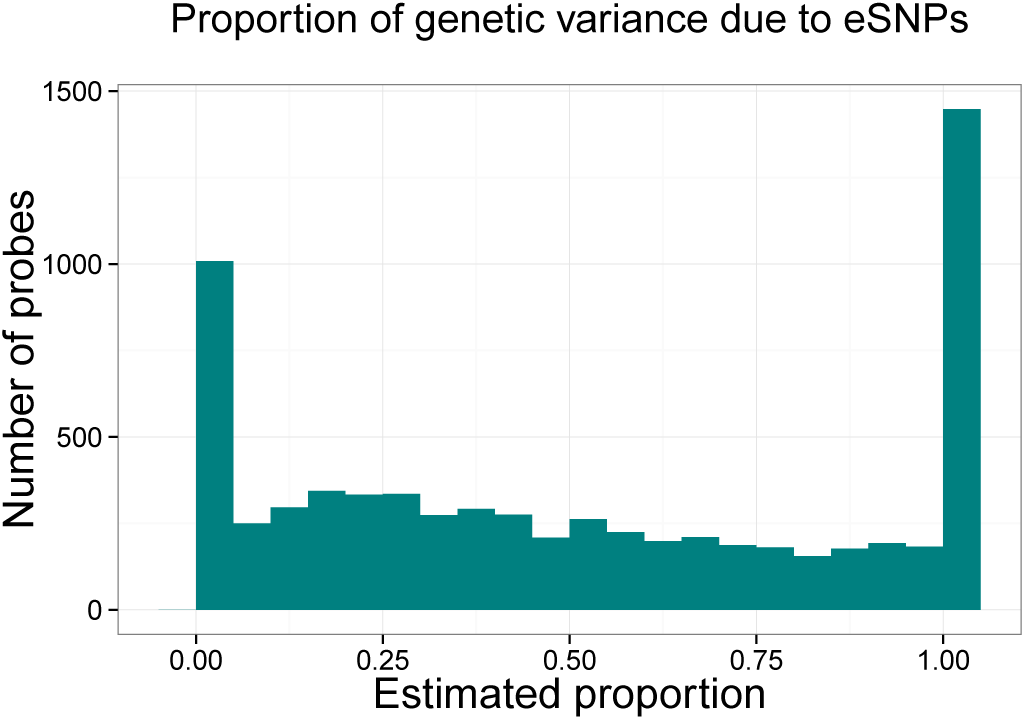
Proportion of genetic variance due to eSNPs (estimated as 1 - the ratio of the genetic variance component when eSNPs are included as fixed effects to the genetic variance component when eSNPs are not included) for the 7,036 heritable probes with local or distal associations, assuming values less than 0 (12%) are exactly 0.

To understand the number of independent signals represented by the eSNPs, we obtained the results from model selection using eSNPs as the pool of possible predictors, focusing again on the set of 7,036 significantly heritable probes with associations to any eSNPs. For this set, the median number of eSNPs with significant marginal association was 24 (mean 42.4), with 470 probes (6.7%) associated to only one eSNP. The distribution of the number of eSNPs included in the best multivariate linear model had a median of 2 (mean 3.1), with 5,212 probes (74%) associated to multiple eSNPs. The large discrepancy in the number of associated SNPs underscores the fact that a substantial proportion of the pairwise SNP-probe associations is due to linkage disequilibrium among neighboring SNPs. At the same time, it is interesting that the selected linear model includes multiple SNPs for 74% of the probes considered: this observation can be interpreted as the result of multiple variants with regulatory effects, but also as a sign that the causal variant is not typed and multiple typed SNPs allow a better reconstruction of the associated haplotype.

To gain insight into the explanatory power of the univariate vs. multivariate models, we assessed the percentage of total phenotypic variance explained by the most significantly associated SNP and by the selected multivariate linear model. The distribution of the percentage of variance explained for the most significantly associated SNP (Figure 4) has a median of 3.7%, a bit lower than results from Zeller et al. (2010) (median=7.7%). The median value of r^2^ increases from 3.7% in the univariate model to 7.4% for the best multivariate model (Figure 4). To understand how much heritability was captured by the linear models involving the eSNPs, we also computed the ratio of the percentage of variance explained for the univariate and multivariate models to the probe heritability estimated using the variance components model. This ratio has median 15% for most significantly associated SNP and 30% for the best multivariate model.

**Figure 4:**
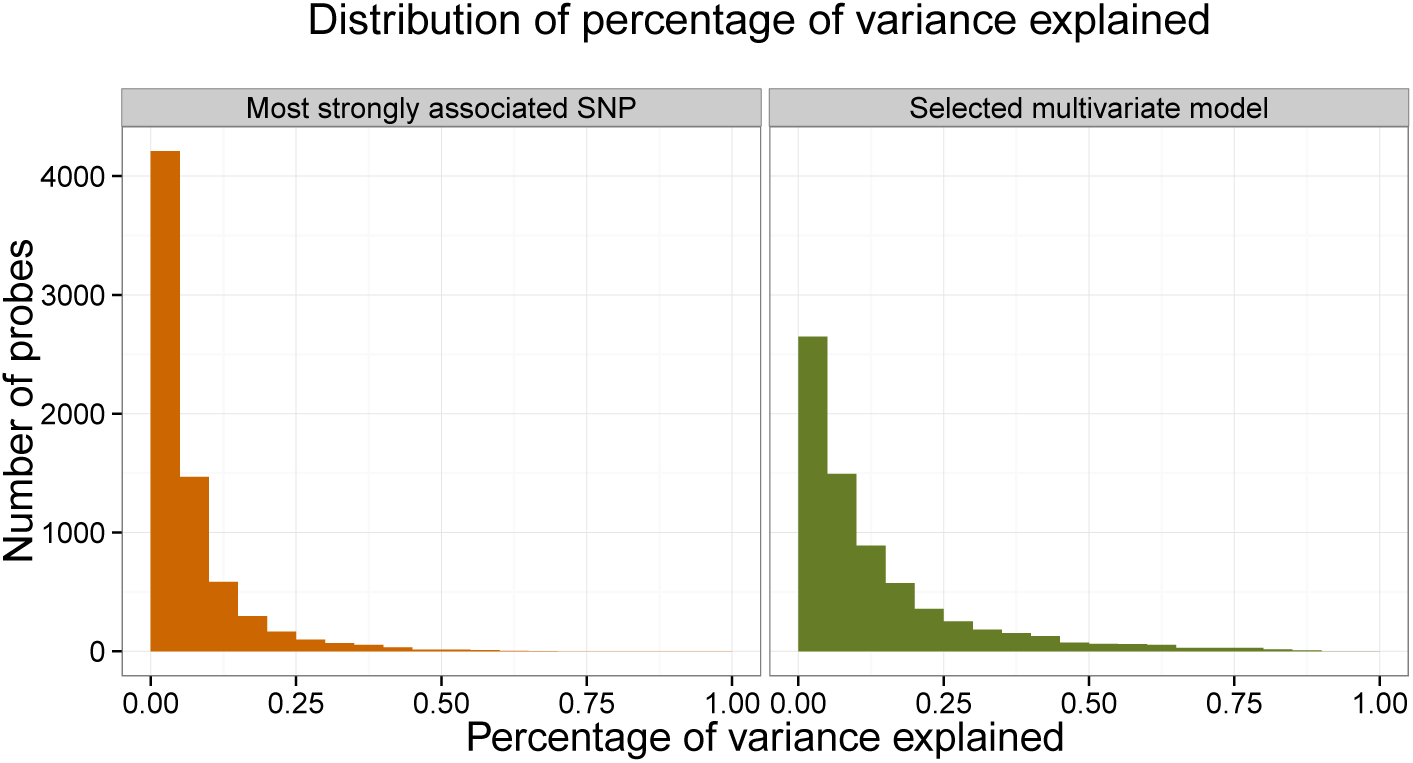
Percentage of variance explained by the most strongly associated eSNP and by the best set of eSNPs selected using multivariate model selection for the 7,036 heritable probes with local or distal eAssociations.

## DISCUSSION

The eQTL study of LCL expression in subjects from extended families segregating for BP1 allows us to tackle questions of general interest as well as possibly identifying regulatory variants specific to this sample. Taking advantage of the pedigree information, we can provide estimates of heritability of the expression traits, as well as compare the results of different estimating procedures, relying on theoretical kinship coefficients or on empirical correlations between observed genotypes. Our results suggest that variation in expression values is heritable and that, at least in samples including related individuals, relying on theoretical kinship coefficients or on realized genotype correlation for estimation of heritability leads to similar results.

Previous studies have obtained a wide range of estimates of the heritability of gene expression, likely due at least in part to the variety of designs that they have employed and tissues evaluated. Our heritability estimates (18% of probes had heritability > 0.2) are larger than those reported by Stranger et al. (2007), who found, in a study of LCL expression in trios from different populations, that 10% and 13% out of 47,294 probes had heritability > 0.2 in Europeans and Yorubans, respectively. In a study of peripheral blood expression in 654 complete twin pairs (1,308 subjects), Wright et al. (2014) report 4.2% of 18.4K genes to be significantly heritable at FDR<5%, and mean heritability of these significantly heritable probes was 0.15.

On the other hand, our heritability estimates are much smaller than those of Göring et al. (2007), who studied lymphocyte expression in large extended families (1,240 subjects in 30 families). They found that 85% of 19,648 probes were significantly heritable at FDR<5%, and median heritability over all probes was 0.23. Grundberg et al. (2012), who studied 856 female twin pairs, obtained similarly high values, estimating that the average heritability of expression in LCL of 23,596 probes is 0.21. Considering only the 17% of probes with cis eQTL at FDR<1%, they found average heritability to be 0.25.

We found that the strategy used for normalization of gene expression had a large impact on the final heritability estimates. Specifically, we observed that normalization of gene expression within pedigrees (following the method described in Kim et al. 2007) inflated estimates of heritability substantially over those obtained using global normalization across all subjects, resulting in values more comparable to those of Göring et al. (2007) and Grundberg et al. (2012). Given that the expression levels of individual genes might be expected to differ across pedigrees, but that global differences are likely due to technical or batch effects, we concluded that the heritability values obtained using within-pedigree normalization were artificially high.

Variance decomposition approaches suggest that 29% of the genetic variance is due to local regulation. In the majority of probes under local regulation in our sample, more than one typed SNP is required to account for expression variation. This finding can be interpreted as the result of heterogeneity, but also could reflect un-typed causal variants that are tracked by more than one typed SNP.

In the effort to control the rate of false discovery among actually reported results (SNPs with apparent regulatory effects), we adopted a hierarchical multiple comparison controlling procedure that is specifically targeted to eQTL studies. It takes into account differences in local and distal regulation, the likelihood that a variant with distal effects might influence the expression of multiple probes, and the dependence between tests for association involving neighboring SNPs. Our major finding is the identification of eSNPs: variants that regulate gene expression. Our results compare favorably with those of more traditional approaches controlling FDR for the entire collection of SNP-probe associations: our local eSNPs are closer to the TSS, and our distal eSNPs regulate more genes.

A question of general interest is how the list of eSNPs we have obtained relates to the genetic underpinnings of the numerous phenotypes available in these pedigrees. Given that the architecture of these traits is more complex than gene expression, and given our limited sample size, gene mapping is more successful for these traits when relying on a linkage approach rather than an association one. The lack of a substantial number of significant SNP associations for these traits makes it impossible to evaluate if eSNPs are enriched in this group. Linkage regions, on the other hand, are wide enough that contrasting the percentage of eSNPs within them and outside them is also rather uninformative. The knowledge we acquired by studying the genetic regulatory network within these pedigrees, instead, can be used to inform our mapping studies: eSNPs might receive a higher prior probability of association, or be assigned a larger portion of the allowed global error rate when using a weighted approach to testing. We will report elsewhere on the results of these investigations.

## SUPPLEMENTARY NOTE

### RNA Extraction and Measurement of Gene Expression

Lymphoblastoid cell lines (LCLs) were established at two sites: RUCDR Infinite Biologics [N=549] and UCLA [N=237]; RNA was extracted from the cell lines at both sites. Peripheral blood mononuclear cells (PBMCs) were separated from venous blood (which had been preserved with an anticoagulating reagent, typically ACD), using either Nycoprep (RUCDR) or Ficoll-Paque PLUS (UCLA) as the separation medium. Both sites used a standard protocol to transform freshly separated or frozen PBMCs, employing media containing Epstein Barr virus (EBV) and the mitogen phytohemagglutinin (PHA). After the culture became established, both sites pelleted cells by centrifugation at 300g for 10 minutes and immediately lyzed them with RLT buffer containing beta-mercaptoethanol, mixed them briefly, and stored them at -80° C until the RNA extraction.

Both RUCDR and UCLA extracted RNA from the cultured cells with an RNeasy 96 kit (Qiagen, Venlo, Netherlands), employing either a Qiagen BioRobot Universal System (RUCDR) or a manual procedure (UCLA) using up to 5×10^5^ cells (RUCDR) or 10^6^ cells (UCLA) as the starting material. To quantify RNA yield, we used a Quant-iT RiboGreen (Invitrogen, Waltham, MA) and measured RNA integrity number, reflective of sample quality, using a TapeStation (Agilent, Santa Clara, CA).

To evaluate gene expression we amplified total RNA (100ng), labeled it using Ambion Total Prep-96 kits (Life Technologies, Grand Island, NY), and hybridized it on Illumina Human HT- 12 v4.0 Expression BeadChips (Illumina Inc., San Diego, CA). Arrays were scanned with an Illumina iScan confocal instrument. Each chip queries the expression of approximately 47,000 probes relative to 31,223 gene targets, as defined by the NCBI reference sequence (RefSeq) database. For these experiments we processed the 786 samples in nine batches (eight batches of between 90–95 samples and one batch of 44 samples).

Data analysis was performed using R (www.r-project.org) and Bioconductor (www.bioconductor.org) packages. Expression values were background corrected, quantile normalized, and log2 transformed using the function necq (Shi et al. 2010). We corrected for the major known batch effects (RUCDR vs. UCLA LCL construction and RNA extraction) using ComBat (Johnson et al. 2007), including BP1 diagnosis, sex, and pedigree IDs as covariates.

### Gene expression quality control

We filtered out 12,834 probes from the initial set of 47,009 probes because of not aligning to hg19 (n=522), not aligning uniquely (1,622), having one or more mismatches in the probe sequence (1,509), or spanning across one or more SNPs in dbSNP 137 or 138 (6,040). After these probes were removed, there were 99 genes interrogated by more than 1 probe where at least one of the probes involved mapped to different locations. Of the mismatching probes, the set with multiple alignments were filtered out (114). In addition, all probes were filtered out for the 12 genes where the probes that map to different chromosomes did not have multiple alignments (31). Finally, 3,141 probes were filtered because they were not detected in at least one sample with an Illumina detection threshold <0.05, leaving 34,030 probes for analysis.

### Genotype quality control and filtering

Genotype data were obtained for 856 subjects. A total of 2,026,257 SNPs were polymorphic and passed all QC procedures. SNPs were excluded in the QC process due to discordance among the three batches in replicated individuals (8,280 SNPs), missing >5% of data (97,158 SNPs), gross violation of Hardy-Weinberg equilibrium (HWE, 79 SNPs), and presence of >4 Mendel errors among fully typed trios (2,976 SNPs). After excluding markers with >4 Mendel errors, the Mendel error rate among fully typed trios was 0.01%, and all further sporadic errors were set to missing in the entire trio. All allele frequency calculations, calculations of HWE, and estimates of LD were performed using only unrelated (founder) individuals. Association analyses used 1,024,051 autosomal SNPs with MAF>10%.

### Screening of subjects

Eighteen subjects were excluded from final analysis because of sample mix-up/contamination (12 subjects) or because they were married-in individuals whose children were not recruited into the study (6 subjects). Among the 838 subjects that passed SNP QC, genotyping completeness was good, averaging 99.78%. Only one subject had genotyping completeness <95%; as they were BP1 and they were missing only 5.7% of genotypes, we retained them for analysis. There were 786 individuals with both genotype and gene expression data (193 BP1 and 593 non-BP1).

